# A Golgi-bypass secretion mechanism for proTGFα involving TMED9 and GRASP65

**DOI:** 10.64898/2026.01.12.698993

**Authors:** Susanne S. Steigleder, Isabel Döring, Julian Nüchel, Marius K. Lemberg

## Abstract

We recently demonstrated that the endoplasmic reticulum export of proTGFα is tightly gated, and that proteostasis factors, including the intramembrane protease RHBDL4, can promote its release by an unknown mechanism. While investigating a potential link between RHBDL4-regulated p24/TMED family proteins and proTGFα trafficking, we uncovered an RHBDL4-independent pathway in which the cargo receptor TMED9 induces unconventional protein secretion (UcPS) of ectopically expressed proTGFα. This pathway selectively requires GRASP65, but not the closely related GRASP55, and depends on components of the autophagic machinery and the ESCRT-associated protein ALIX, consistent with a Golgi-bypass secretion route. Although largely based on overexpression systems, our findings identify a previously unrecognized UcPS mechanism and provide insight into alternative protein trafficking pathways that may be exploited under conditions of proteostasis stress and in pathological contexts such as cancer and inflammation.

## Introduction

It has been a long-standing dogma that proteins have to follow the conventional secretory pathway in cells to be trafficked to the extracellular space or the plasma membrane. This pathway involves co-translational targeting and insertion of cargo molecules into the endoplasmic reticulum (ER) followed by trafficking to the Golgi apparatus. However, over the past years, several unconventional protein secretion (UcPS) pathways have been identified that omit the key organelles of the classical secretory route. Generally, UcPS can be classified into four types. Proteins without signal peptides can be secreted through plasma membrane pores (type I; e.g. FGF2), via ABC transporters (type II), or through secretory organelles as endosomes or autophagosomes (type III) (Duran et al., 2010, Nüchel et al., 2018, Chiritoiu et al., 2018, Sparn et al., 2022). Type IV is mechanistically distinct in that it involves proteins with canonical ER-targeting signal peptides and transmembrane domains that are first synthesized in the ER before reaching the cell surface or extracellular space by bypassing the Golgi apparatus. This group encompasses multiple different pathways, including pathways involving parts of the autophagy machinery (Yoo et al., 2002, Schotman et al., 2008, Gee et al., 2011, Lerche et al., 2025).

One of the best-characterized type IV UcPS cargos is the F508-deletion mutant of the cystic fibrosis transmembrane conductance regulator (CFTRΔF508). This mutant can be secreted unconventionally in response to cellular stress, ER-to-Golgi blockade, or overexpression of the Golgi reassembly-stacking protein (GRASP) 55 or its closely related homolog GRASP65 (Shorter et al., 1999, Gee et al., 2011). In particular, GRASP55 has been implicated in UcPS of CFTRΔF508 and a growing number of additional cargos across multiple contexts (Nüchel et al., 2018, Nüchel et al., 2021, Ahat et al., 2022, Noh et al., 2022). In contrast, the contribution of GRASP65 to type IV UcPS is less well defined. While initial studies have implicated both GRASP55 and GRASP65 in unconventional trafficking of CFTRΔF508 and α5β1integrins, the extent to which GRASP65 is involved in cargo-specific UcPS, and whether it functions independently of GRASP55, remains unclear (Gee et al., 2011, Lerche et al., 2025).

Originally, GRASPs were described as tethering factors required for maintaining Golgi stacks, but in recent years they have been shown to participate in many other processes such as Golgi enzyme recycling (Pothukuchi et al., 2021, Akaaboune et al., 2025, Rabouille and Linstedt, 2016). Both GRASPs are N-terminally myristoylated and thereby tethered to Golgi membranes, but under certain conditions they have also been detected in other subcellular locations depending on posttranslational modifications (Kim et al., 2016, Kortvely et al., 2016, Zhang et al., 2018, Nüchel et al., 2021). Recently, members of the p24/TMED cargo receptor family have also been linked to type III and type IV UcPS (Zhang et al., 2020, Park et al., 2022, Zheng et al., 2025). In vertebrates, this cargo receptor protein family comprises nine members that have a broad and partially overlapping cargo repertoire (reviewed in Pastor-Cantizano et al., 2016). TMED10 has been shown to mediate secretion of a variety of proteins lacking signal peptides by a type III UcPS pathway, in a process that is thought to involve formation of an oligomeric TMED10 pore at the plasma membrane (Zhang et al., 2020). This concept was recently expanded by the demonstration that several additional p24 family members also contribute to type III UcPS (Zheng et al., 2025). In addition, TMED2, TMED3, TMED9, and TMED10 have been shown to form a complex that promotes type IV UcPS of CFTRΔF508 and H723R-pendrin in a process that is at least partially dependent on GRASP55 (Park et al., 2022).

Transforming growth factor α (TGFα) is synthesized as a pro-form as a type I membrane protein that is cleaved at two sites by ADAM metalloproteases, sequentially releasing its propeptide and the bioactive TGFα ligand - in a process known as shedding (Peschon et al., 1998). The mature TGFα activates the epidermal growth factor (EGF) receptor, promoting cell proliferation and development, and has been associated with cancer (reviewed in Singh and Coffey, 2014). However, the ER-to-Golgi trafficking of proTGFα remains only partially understood. The p24 protein TMED9 and the Cornichon cargo receptors have both been linked to proTGFα trafficking (Zhang and Schekman, 2016, Mishra et al., 2019). Increased secretion of TGFα has previously been linked to bad prognosis in various cancer types (Ma et al., 2024, Yang et al., 2025). Intriguingly, proTGFα interacts with GRASP55 via its C-terminus, and disrupting this interaction reduces its cell surface localization (Kuo et al., 2000), suggesting a role of GRASP55 in its conventional trafficking. More recently, we showed that ectopically expressed proTGFα is predominantly targeted for ER-associated degradation (ERAD), but the intramembrane protease RHBDL4 can promote ER export leading to a strong increase in its secretion, including a microvesicle-based pathway (Wunderle et al. 2016). In this study, we show that TMED9 overexpression induces UcPS of proTGFα independently of RHBDL4. Moreover, GRASP65, but not GRASP55 acts in the same pathway as TMED9 and promotes unconventional transport of proTGFα. Mechanistically, we can show that this UcPS mechanism depends on autophagy components such as ATG5/ATG7 and involves the ESCRT-associated protein ALIX.

## Results

### TMED9 overexpression induces Golgi-bypass UcPS of proTGFα

Ectopically expressed proTGFα is largely retained in the ER and targeted for ERAD. We previously showed that the intramembrane protease RHBDL4 can promote ER export and microvesicle-mediated secretion of proTGFα through a mechanism that remains unresolved (Wunderle et al., 2016). In light of our recent finding that RHBDL4 cleaves selected p24/TMED cargo receptors (Knopf et al., 2024), we asked whether members of the TMED family participate in this proTGFα-tuning mechanism. To test this hypothesis, we co-expressed the two most efficiently cleaved p24 proteins, namely TMED3 and TMED7, as well as the non-substrate TMED9 (Knopf et al., 2024), with FLAG-proTGFα (Fig. 1A, B). To prevent proTGFα shedding by ADAM proteases, cells were treated with the metalloprotease inhibitor BB94. Overexpressed FLAG-proTGFα can be primarily observed in the cell extract as a 28 kDa and a 35 kDa form (Fig. 1B). As observed before (Wunderle et al., 2016), treatment with the deglycosylating enzymes EndoH and PNGaseF revealed, that, in contrast to the 35-kDa form, the glycan of the 28 kDa form was not modified in the Golgi apparatus (Fig. 1C). Hence, the 28-kDa species represents the ER-based form of proTGFα. In contrast, the 35 kDa species is EndoH insensitive, indicating that it has reached or passed the Golgi.

**Fig. 1.**
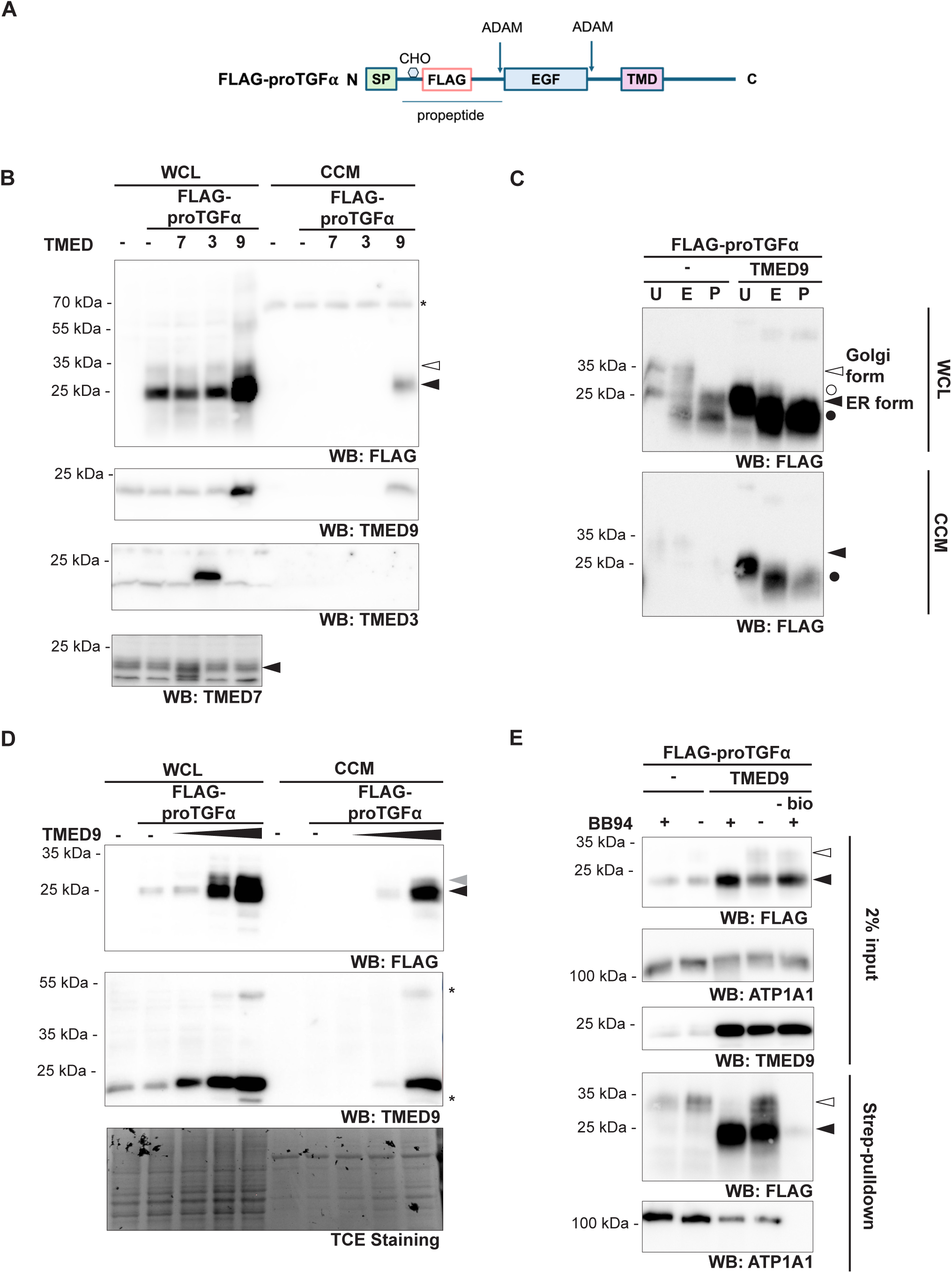
TMED9 overexpression induces intracellular accumulation and UcPS of proTGFα. **(A)** Schematic of FLAG-proTGFα expression construct used. SP, signal peptide, TMD, transmembrane domain. **(B)** Western blot (WB)-based secretion assay using Hek293T cells co-expressing FLAG-proTGFα with either empty vector (-), TMED3, TMED7 or TMED9. FLAG-proTGFα can be observed as a 28-kDa ER form (filled arrow) and a 35-kDa Golgi form (open arrow). TMED9 co-expression promotes intracellular accumulation and secretion of the ER form. Background bands are marked with an asterisk. Cells were treated with BB94 (10 µM). WCL, whole cell lysate, CCM, concentrated conditioned medium. **(C)** WB analysis following no treatment (U) or treatment with EndoH (E) or PNGaseF (P) of Hek293T cells co-expressing FLAG-proTGFα either with an empty vector (-) or TMED9. The TMED9-induced secreted FLAG-proTGFα species (filled arrow) is sensitive to EndoH and PNGaseF (filled circle), whereas the Golgi form (open arrow) is sensitive to PNGaseF (open circle). Cells were treated with BB94 (10 µM). **(D)** Dose-dependent effect of TMED9 on FLAG-proTGFα in HCT116 cells. Increasing amounts of TMED9 (100 ng, 500 ng, 1000 ng) induces intracellular accumulation and secretion of the ER form (filled arrow) and an intermediate form (grey arrow). TCE staining is used as a loading control. Unknown bands are indicated with an asterisk. Cells were treated with BB94 (10 µM). **(E)** WB-based cell surface biotinylation assay in Hek293T cells co-expressing FLAG-proTGFα with either empty vector (-) or TMED9. TMED9 overexpression induces increased localization of the ER form of proTGFα (filled arrow) at the cell surface. A non-biotinylated sample (- bio) serves as negative control. ATP1A1 signal serves as positive control. Where indicated, cells were treated with BB94 (10 µM).

While TMED3 and TMED7 expression showed no effect on proTGFα, TMED9 expression led to increased intracellular steady-state level and apparent secretion of the 28-kDa form, which co-migrated with the ER form of proTGFα observed in the cell extract (Fig. 1B). Interestingly, overexpressed TMED9 itself was also detected in the medium fraction, suggesting that it traffics and is co-released in extracellular vesicles together with proTGFα. Consistent with secretion via a Golgi-independent UcPS pathway, the secreted proTGFα form was sensitive to EndoH treatment (Fig. 1C), suggesting that it had not traversed the Golgi. TMED9 and TGFα were recently both linked to colorectal cancer metastasis (Mishra et al., 2019). Consistently, these effects were not restricted to Hek293T cells but were also observed in the colon cancer cell line HCT116 in a dose-dependent manner (Fig. 1D). In this context, an additional 30-kDa intermediate species of proTGFα was apparent, likely corresponding to a partially complex glycosylated form.

Since proTGFα is known to also localize at the plasma membrane (Teixido et al., 1990), we wondered whether ectopic TMED9 expression might also induce unconventional trafficking to the cell surface. Supporting this idea, cell surface biotinylation revealed that ectopic TMED9 expression promoted cell surface localization of the 28-kDa proTGFα form, which co-migrated with the ER-based species (Fig. 1E). Surprisingly, this effect was insensitive to BB94 treatment, indicating that the surface localized pool of proTGFα was inaccessible to ADAM sheddases. In contrast, and consistent with earlier reports (Wunderle et al., 2016), the Golgi-modified 35-kDa form was only detected when ADAM proteases were inhibited (Fig. 1E). These findings suggest that ectopically expressed TMED9 induces UcPS-mediated secretion, leading to cell surface localization of proTGFα. However, comparison of the glycosylation pattern argues that this effect is likely independent of the RHBDL4-dependent pathway, which, as observed previously, enhances the 35-kDa Golgi-modified form of proTGFα (Fig. S1) (Wunderle et al., 2016).

Next, we investigated which features are responsible for the TMED9-mediated effect. TMED9 contains a di-phenylalanine motif in its C-terminus to interact with the COPII machinery (Nufer et al., 2002). To test whether COPII interaction by TMED9 is required to induce UcPS of proTGFα, we abrogated the motif by replacing it with alanine (TMED9-AA) (Fig. S2A). Consistent with the motif being required for TMED9-mediated UcPS induction of proTGFα, TMED9-AA expression neither led to intracellular accumulation of the ER form of proTGFα nor induced its secretion (Fig. S2B). However, TMED9-AA was also not detected in the medium fraction (Fig. S2B), and the entire TMED9-AA pool was sensitive to EndoH, indicating that it is retained in the ER (Fig. S2C). Hence, the lack of UcPS upon TMED9-AA expression could result either from its inability to interact with the COPII machinery or from a failure to exit the ER.

### GRASP65 but not GRASP55 overexpression phenocopies TMED9-induced UcPS

Given that GRASP55 has been linked to TMED-induced UcPS of CFTRΔF508 (Park et al., 2022), we wondered whether it is involved in TMED9-induced UcPS of proTGFα. GRASP55 and its closely related homolog GRASP65 were co-expressed as myc-tagged expression constructs in different amounts with proTGFα (Fig. 2A). Notably, GRASP65 expression induced a phenotype similar to that of TMED9, with dose-dependent intracellular accumulation of the ER-form of proTGFα, alongside the 30-kDa intermediate form previously observed upon TMED9 expression in HCT116 cells (Fig. 1D), both of which were secreted (Fig. 2A). Upon GRASP55 expression, a similar but markedly less pronounced effect was observed (Fig. 2A). Of note, both ectopically expressed GRASP65 and GRASP55 were detected in the medium fraction. Similar observations were made in HCT116 cells (Fig. 2B). Similar to TMED9, GRASP65 induced unconventional proTGFα secretion, as the secreted form was sensitive to EndoH treatment (Fig. 2C). Notably, GRASP65 expression led to increased plasma membrane localization of both the ER- and Golgi-associated forms of proTGFα, which were not sensitive to ADAM-dependent shedding, as their levels were not reduced in absence of BB94 treatment (Fig. 2D). Our results demonstrate that GRASP65, and to a minor extent GRASP55, induce UcPS of proTGFα in a manner similar to TMED9.

**Fig. 2.**
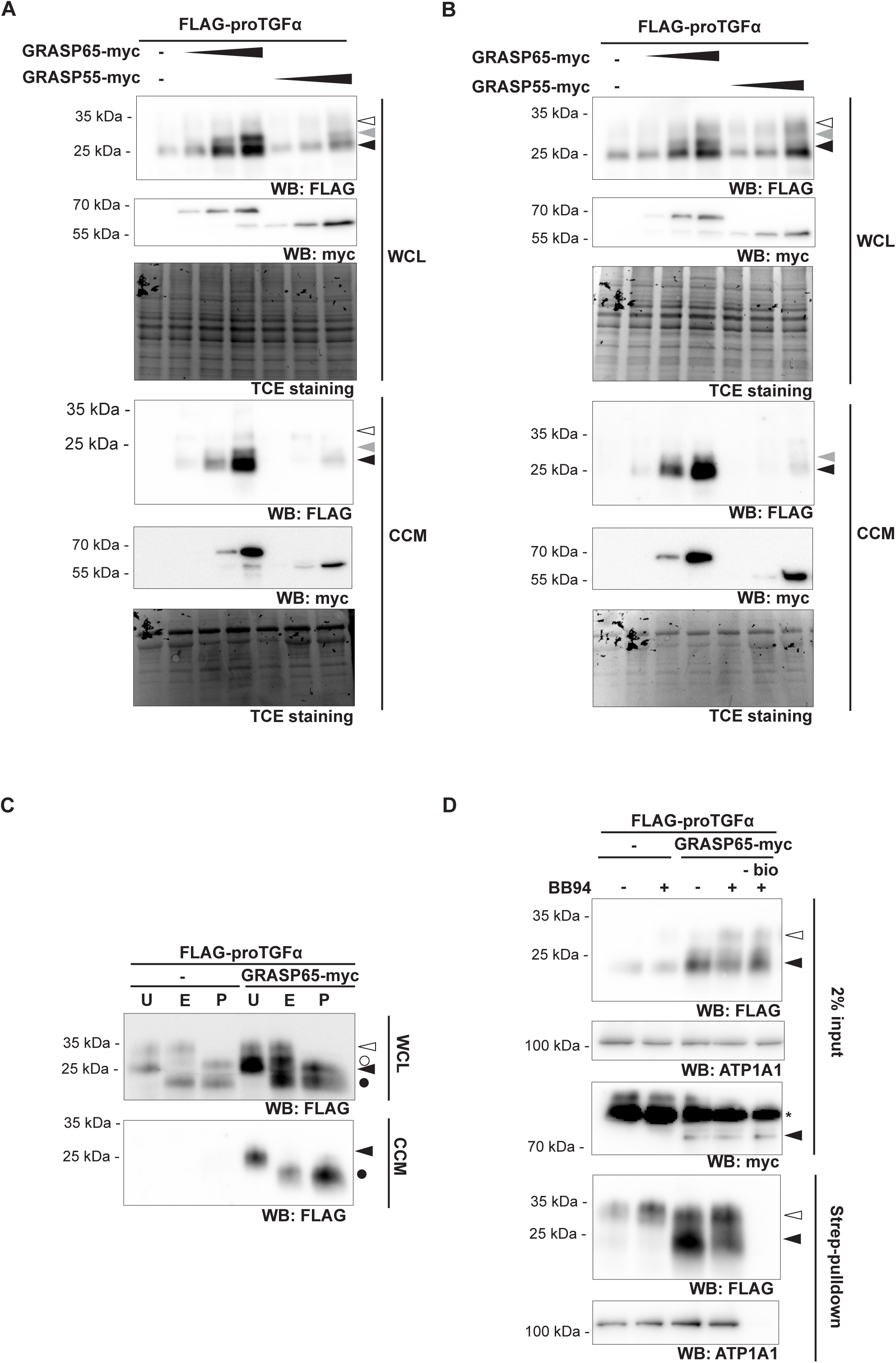
Ectopic expression of GRASP65 phenocopies TMED9-mediated UcPS of proTGFα. **(A)** Western blot (WB)-based secretion assay of Hek293T cells co-expressing FLAG-proTGFα with different amounts of either empty vector (-), GRASP55-myc or GRASP65-myc (100 ng, 500 ng, 1000 ng). GRASP65-myc, but not GRASP55-myc, induces dose-dependent intracellular accumulation and secretion of the ER form (filled arrow) and an intermediate form (grey arrow) of proTGFα, while the Golgi form (open arrow) remains unchanged. TCE staining is used as a loading control. Cells were treated with BB94 (10 µM). **(B)** Same experiment as in (A), performed in HCT116 cells, yielding comparable results. **(C)** WB analysis following no treatment (U) or treatment with EndoH (E) or PNGaseF (P) of Hek293T cells co-expressing FLAG-proTGFα with empty vector (-) or GRASP65-myc. The GRASP65-induced secreted FLAG-proTGFα species (filled arrow) is sensitive to EndoH (filled circle), while the Golgi form (open arrow) is sensitive to PNGase (open circle). Cells were treated with BB94 (10 µM). **(D)** Cell surface biotinylation assay in Hek293T cells co-expressing FLAG-proTGFα with either empty vector (-) or GRASP65-myc. GRASP65 expression increases surface exposure of the 28-kDa ER form of proTGFα (filled arrow). A non-biotinylated sample (- bio) serves as negative control. ATP1A1 signal serves as positive control. Where indicated, cells were treated with BB94 (10 µM).

### GRASP65-mediated UcPS depends on its SPR domain and the proTGFα PDZ-binding domain

Both GRASP proteins share the same domain structure, composed of two N-terminal PDZ domains of high sequence identity, followed by a more diverse C-terminal serine-proline rich (SPR) domain (Fig. 3A) (Shorter et al., 1999, Wang et al., 2005). To test which domain of GRASP65 is required for the induction of proTGFα UcPS, we co-expressed proTGFα together with chimeric GRASP55/GRASP65 mutants (Nüchel et al., 2021) (Fig. 3A). Notably, only the mutant containing the GRASP65-SPR domain induced UcPS of proTGFα to a similar extent as wild-type GRASP65 (Fig. 3B).

**Fig. 3.**
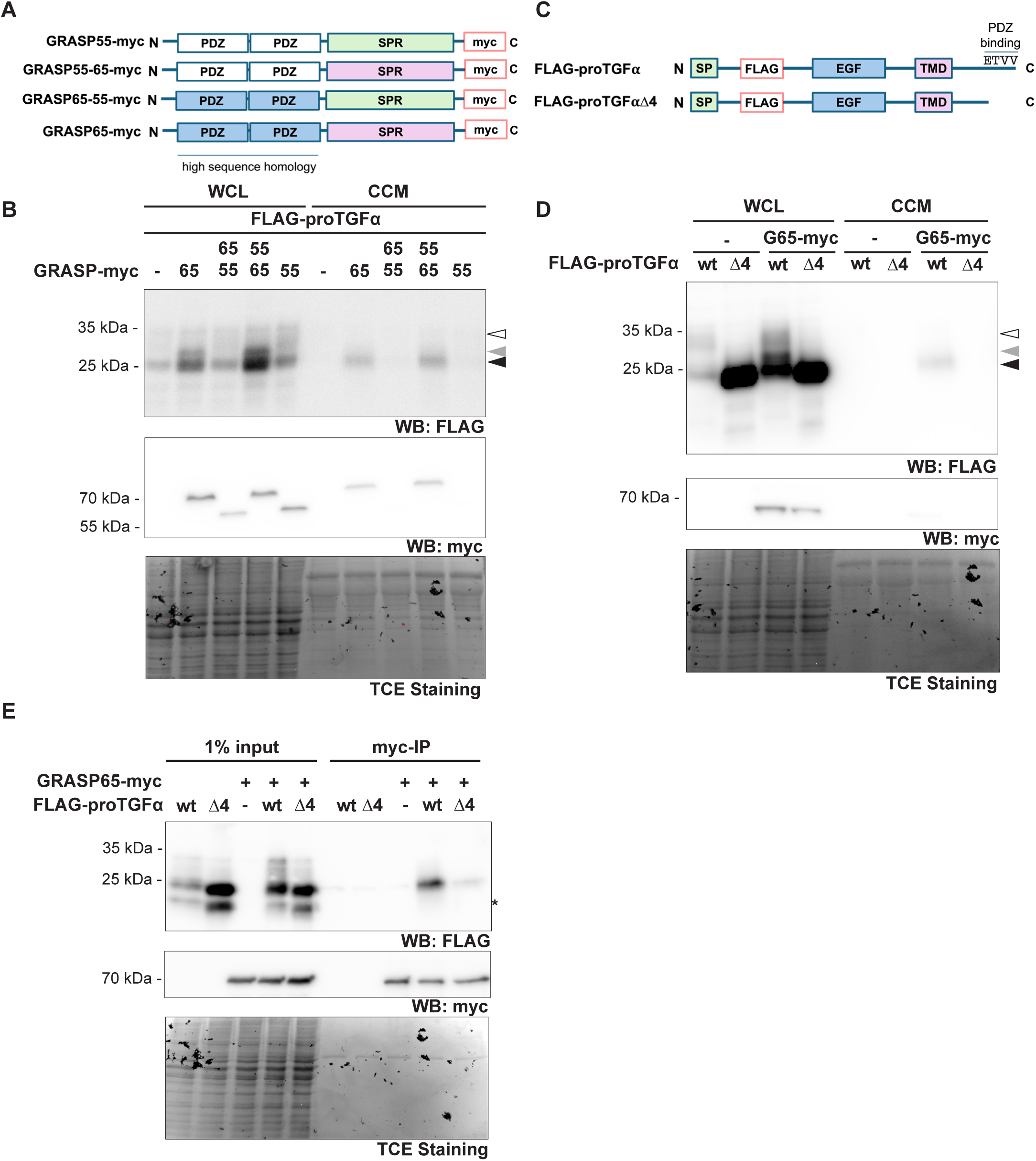
GRASP65-mediated UcPS depends on its SPR domain and the proTGFα PDZ-binding domain. **(A)** Schematic representation of GRASP55/GRASP65 chimeric constructs with exchanged PDZ-domains and SPR-domains. **(B)** Western blot (WB)-based secretion assay using Hek293T cells co-expressing FLAG-proTGFα with either empty vector (-), GRASP55-myc (55), GRASP65-myc (65) or the chimeric mutants (55-65, 65-55). Only the 55-65 chimera phenocopies GRASP65-myc, indicating a requirement for the GRASP65 SPR domain. TCE staining is used as a loading control. Cells were treated with BB94 (10 µM). **(C)** Schematic of FLAG-proTGFα and the Δ4 mutant, which lacks the PDZ-binding domain (last four amino acids). SP, signal peptide, TMD, transmembrane domain. **(D)** WB-based secretion assay using Hek293T cells co-expressing either wild-type (wt) or Δ4 FLAG-proTGFα with empty vector (-) or GRASP65 (G65)-myc. Deletion of the PDZ-binding motif abolishes GRASP65-induced UcPS. TCE staining is used as a loading control. Cells were treated with BB94 (10 µM). **(E)** Myc-specific immunoprecipitation (IP) from Hek293T cells co-expressing GRASP65-myc with either empty vector (-), wt FLAG-proTGFα, or FLAG-proTGFαΔ4. WB analysis shows strongly reduced interaction between GRASP65-myc and the Δ4 mutant. Unknown bands are indicated with an asterisk. TCE staining serves as loading control.

To further investigate the potential interaction between GRASP65 and proTGFα, we deleted the PDZ-binding motif in the proTGFα cytosolic tail (proTGFαΔ4) (Fig. 3C) (Briley et al., 1997, Singh and Coffey, 2014), which may bind to PDZ-domains of GRASP65. Consistently, co-expression of GRASP65 with proTGFαΔ4 failed to induce UcPS (Fig. 3D) and interaction between GRASP65 and proTGFαΔ4 is profoundly reduced as assessed by co-immunoprecipitation (Fig. 3E). These findings suggest that the SPR domain of GRASP65 is required to mediate efficient UcPS of proTGFα, while the interaction between the two proteins occurs via the PDZ-binding motif of proTGFα and the PDZ domains of GRASP65, in agreement with previous yeast-two-hybrid data (Barr et al., 2001).

### TMED9 and GRASP65 interact and function in the same UcPS pathway

The similarity of the effects induced by TMED9 and GRASP65 overexpression on proTGFα suggest that they act in the same pathway. Consistent with this notion, TMED9 knockdown markedly reduced secretion of the 28-kDa proTGFα species and, notably, also decreased the amount of secreted GRASP65 (Fig. 4A). We therefore asked whether TMED9 and GRASP65 physically interact. Because such interactions may be transient, we performed immunoprecipitation using the cleavable crosslinker dithiobis(succinimidylpropionate) (DSP) to stabilize weak or short-lived associations (Fig. 4B). Under native conditions, only a small fraction of TMED9 co-purified with GRASP65, whereas DSP crosslinking revealed a robust interaction (Fig. 4B). It remains to be determined which domain of GRASP65 is required for this interaction. We further tested whether proTGFα influences the TMED9-GRASP65 interaction and observed no major changes, although proTGFα co-purified with GRASP65 both in the presence and absence of DSP (Fig. 4B) consistent with our earlier findings (Fig. 3E). Together, these data indicate that TMED9 and GRASP65 act within the same UcPS pathway and physically associate during proTGFα UcPS; however, the precise step at which this interaction occurs, and whether it reflects a direct to complex-mediated contact, remains unresolved.

**Fig. 4.**
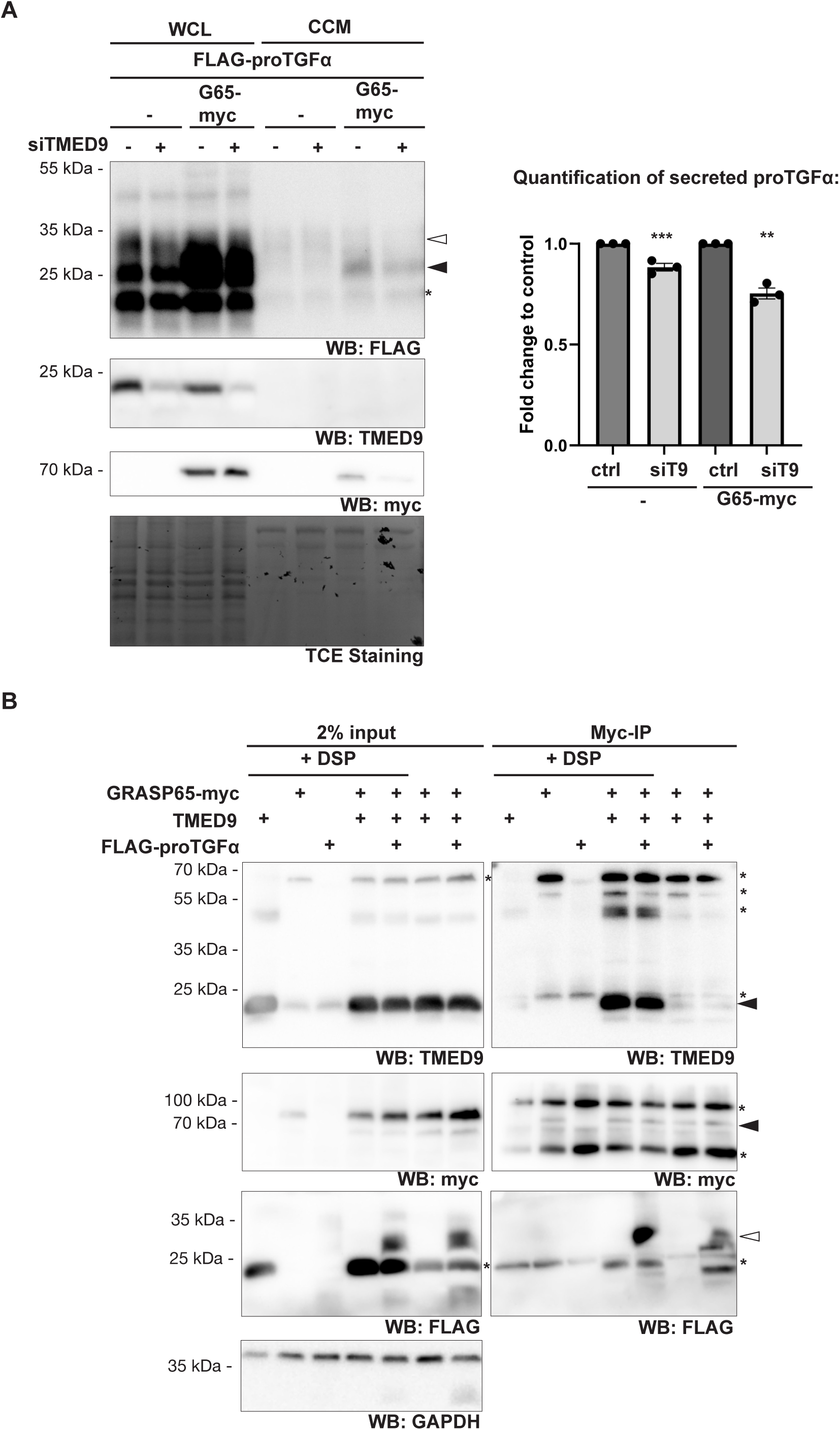
TMED9 and GRASP65 interact and function in the same UcPS pathway. **(A)** Western blot (WB)-based secretion assay using Hek293T cells co-expressing FLAG-proTGFα with empty vector (-) or GRASP65-myc (G65-myc), combined with control or TMED9-specific siRNA (siTMED9). TMED9 knockdown reduces secretion of the proTGFα ER-form (filled arrow) (means ± SEM, n = 3, **p < 0.01, ***p < 0.001, unpaired two-sided Student’s t-test). Unknown bands are indicated with an asterisk. TCE staining is used as a loading control. Cells were treated with BB94 (10 µM). **(B)** Co-immunoprecipitation (IP) analysis of Hek293T cells expressing either GRASP65-myc, TMED9, and/or FLAG-proTGFα. Lysates were prepared with Triton X-100 and subjected to myc-specific IP. A robust TMED9-GRASP65 interaction is detected upon crosslinker DSP (filled arrow) and is not affected by FLAG-proTGFα co-expression. Background bands are indicated by an asterisk. GAPDH serves as loading control. Cells were treated with BB94 (10 µM).

### TMED9- and GRASP65-induced UcPS of proTGFα requires ALIX

One established UcPS route for membrane proteins involves secretory autophagy. We therefore asked whether the autophagy machinery contributes to TMED9- and GRASP65-induced UcPS of proTGFα. To achieve efficient inhibition of autophagy, we simultaneously ablated the two core components ATG5 and ATG7 (Galluzzi and Green, 2019). This resulted in a significant reduction of TMED9- or GRASP65-induced UcPS of proTGFα (Fig. 5A). Notably, intracellular accumulation of proTGFα was also decreased, likely reflecting reduced UcPS activity upon TMED9 or GRASP65 expression under autophagy-deficient conditions (Fig. 5A). Together, these findings indicate that secretory autophagy contributes to the UcPS of proTGFα.

**Fig. 5.**
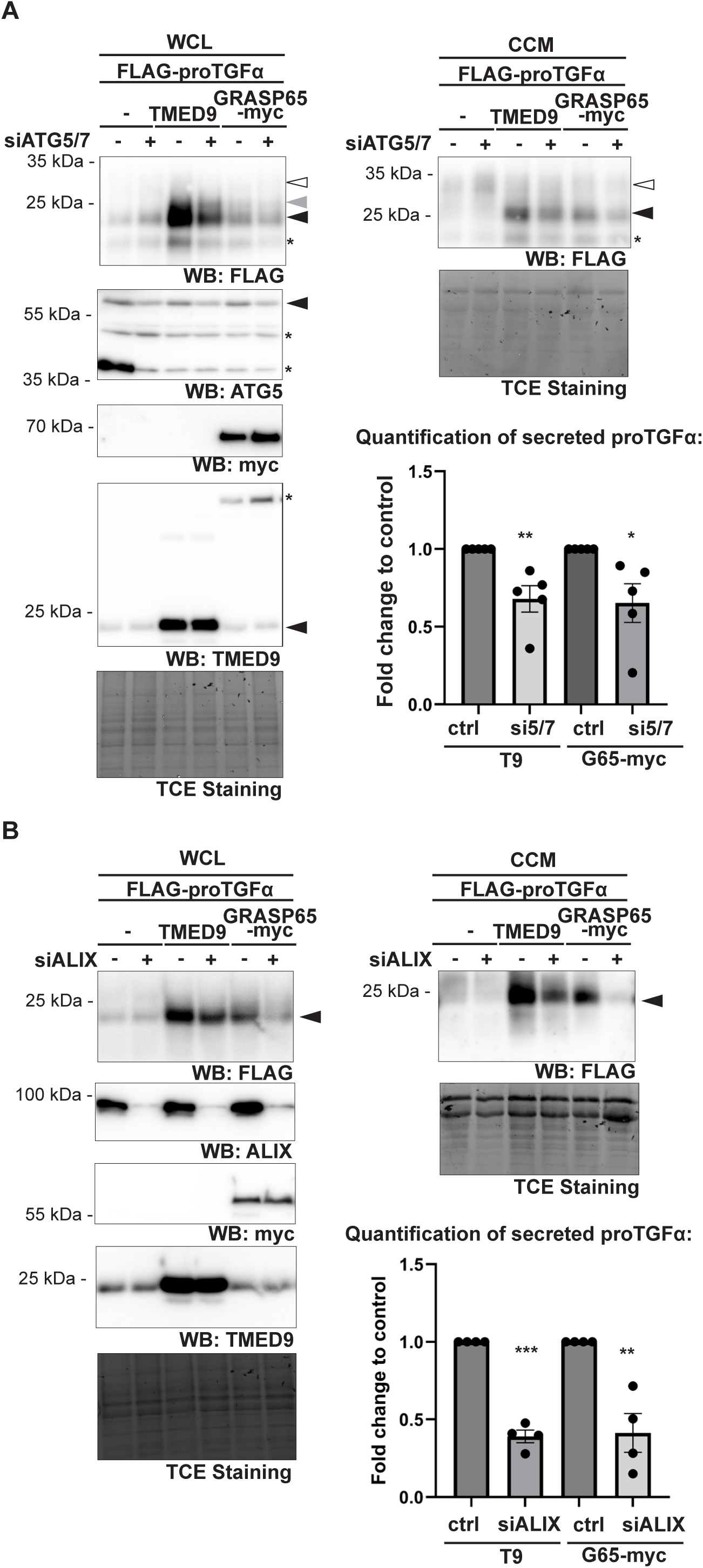
TMED9- and GRASP65-induced UcPS of proTGFα requires ALIX. **(A)** Western blot (WB)-based secretion assay using Hek293T cells co-expressing FLAG-proTGFα with empty vector (-), TMED9, or GRASP65-myc, combined with control or ATG5/ATG7 double knockdown (siATG5/7). Depletion of ATG5 and ATG7 significantly reduces TMED9 (T9)- and GRASP65-myc(G65-myc)-mediated UcPS of proTGFα (means ± SEM, n = 3, *p < 0.05, **p < 0.01, unpaired two-sided Student’s t-test). Unknown bands are indicated with an asterisk. TCE staining is used as a loading control. Cells were treated with BB94 (10 µM). **(B)** Same assay as in (A), performed with ALIX-specific siRNA (siALIX). ALIX depletion markedly reduces TMED9- and GRASP65-myc-induced UcPS of proTGFα (means ± SEM, n = 3, **p < 0.01, ***p < 0.001, unpaired two-sided Student’s t-test). TCE staining is used as a loading control. Cells were treated with BB94 (10 µM).

Recently, it was demonstrated that ER exit sites (ERES) can be degraded by autophagy and that this process is mediated by components of the ESCRT machinery (Liao et al., 2024). This degradation occurs under nutrient starvation, which is a condition that also typically induces UcPS pathways via the mTORC1-GRASP55 axis (Nüchel et al., 2021). This raises the possibility that this mechanism may be linked to a form of secretory autophagy. In this context, the ESCRT-associated protein ALIX was identified as a key factor (Liao et al., 2024). ALIX binds to ESCRT-III CHMP4 proteins and the ESCRT-I subunit TSG101, thereby acting as a scaffold for ESCRT assembly (Strack et al., 2003, Katoh et al., 2004, Morita et al., 2007, McCullogh et al., 2008). However, its role in multivesicular body (MVB) formation appears to be cell type- and cargo-specific (Cabezas al., 2005, Baietti, et al., 2012, Colombo et al., 2013). Consistent with a functional contribution, knockdown of ALIX strongly reduced TMED9- and GRASP65-induced UcPS of proTGFα (Fig. 5B). This suggests that, in addition to its role in degradative autophagy of ERESs, ALIX may also participate in UcPS of proTGFα. Alternatively, UcPS of proTGFα may rely on ALIX-dependent MVB biogenesis.

## Discussion

Here, we identify a previously unrecognized UcPS pathway involving TMED9 and GRASP65, using ectopically expressed proTGFα as a model cargo. This work extends our previous observation that secretion of surplus proTGFα can be enhanced by the intramembrane protease RHBDL4, leading to its release in microvesicles (Wunderle et al., 2016). Our current findings indicate that the TMED9- and GRASP65-dependent pathway bypasses the Golgi apparatus and is mediated by secretory autophagy independent of RHBDL4. Notably, several cancer types exhibit elevated expression and secretion of proTGFα, raising the question whether endogenous proTGFα, particularly in high expressing cancer cell lines, is subject to TMED9- or GRASP65-mediated UcPS under physiological or stress conditions.

### Mechanistic model for proTGFα UcPS

Under overexpression conditions, newly synthesized proTGFα exceeds the tightly regulated ER export capacity and is therefore predominantly eliminated by ERAD (Horiuchi et al., 2007, Wunderle et al., 2016) (Fig. 6). Only a minor fraction reaches the cell surface via the conventional Golgi-dependent route, where it can be shed by ADAM metalloproteases or released in microvesicles (Peschon et al., 1998, Wunderle et al., 2016). Co-expression of TMED9 or GRASP65 actively redirects proTGFα away from proteasomal degradation and promotes its transport to the cell surface and secretion through a Golgi-bypass UcPS pathway (Fig. 6).

**Fig. 6.**
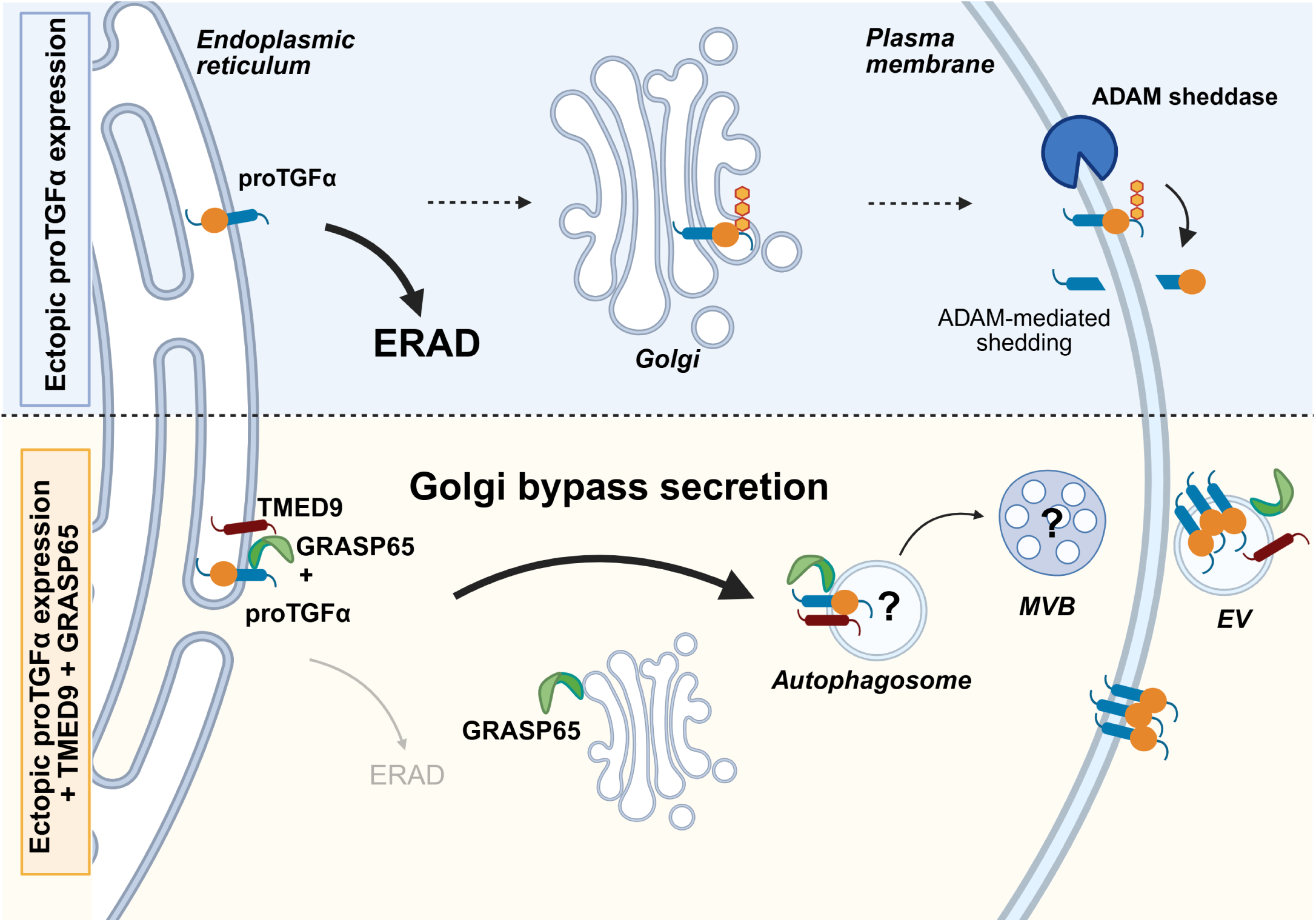
Proposed mechanism of TMED9- and GRASP65-induced UcPS of proTGFα. Under conditions of overexpression, newly synthesized proTGFα is predominantly degraded by ERAD (Wunderle et al., 2016), whereas only a minor fraction enters the conventional secretory pathway and is shed at the plasma membrane. Co-expression of TMED9 or GRASP65 diverts proTGFα away from degradation, leading to its accumulation in pre-Golgi compartments and possibly the cis-Golgi. Within these compartments, proTGFα interacts with GRASP65 and enters a Golgi-bypass pathway mediated by secretory autophagy potentially involving incorporation of proTGFα into autophagosomes that can fuse with MVBs) prior to secretion. ProTGFα is subsequently released into the extracellular space within extracellular vesicles (EV). Created with https://biorender.com.

TMED9 and GRASP65 are co-secreted with proTGFα, suggesting formation of a joint trafficking complex, either directly or indirectly. Consistent with this model, TMED9 and proTGFα have previously been detected in COPII vesicles (Cao et al., 2008), and both we and others observe a physical interaction between GRASP-proteins and proTGFα (Barr et al., 2001). The proTGFα PDZ-binding motif is required for GRASP65-induced UcPS, likely reflecting a recognition mechanism analogous to the known interaction between proTGFα and GRASP55 in conventional secretion (Kuo et al., 2000). A defining feature of both pathways is the requirement of the GRASP-specific SPR domains (Kuo et al., 2000). Given that phosphorylation of the GRASP55 SPR domain is essential for UcPS-induction of other cargoes (Gee et al., 2011, Nüchel et al., 2021), analogous regulation of GRASP65 may promote UcPS of proTGFα.

TMED9 and GRASP65 appear to act within the same pathway but may associate only transiently, potentially during assembly of a cargo-containing trafficking intermediate. Depletion of ATG5 and ATG7 markedly reduced proTGFα secretion, indicating that this unconventional route requires components of the autophagic machinery and suggesting incorporation of proTGFα into secretory autophagosomes (Fig. 6). This is consistent with previous links between GRASP proteins and autophagy: GRASP65 contains several LC3-interacting regions, and the homologous motifs in GRASP55 are required for LC3-binding (Zhang et al., 2018, Nüchel et al., 2018) and for UcPS of TGFβ1 (Nüchel et al., 2018). TMED9 has likewise been implicated in autophagosome biogenesis via a motif adjacent to its COPII-binding site (Li et al., 2022).

Secretory autophagy frequently intersects with the endosomal system, and autophagosomes can fuse with MVBs in an ESCRT-dependent manner (Hessvik and Llorente, 2018). Accordingly, depletion of the ESCRT-associated protein ALIX attenuated proTGFα UcPS, suggesting a role for MVB-related intermediates. Alternatively, recent evidence indicates that ERES can be degraded by autophagy in an ATG7- and ALIX-dependent process (Liao et al., 2024), which may represent an additional or parallel, yet undiscovered route for secretion. Thus, while our data support a role for secretory autophagy in proTGFα UcPS, the precise molecular mechanism remains to be defined.

### Potential physiological relevance and link to CFTRΔF508 UcPS

Although this study focuses on ectopically expressed proTGFα, it is notable that under these conditions proTGFα, similar to CFTRΔF508, is predominantly targeted for ERAD unless UcPS is engaged (Gee et al., 2011; Wunderle et al., 2016; Noh et al., 2018). Both cargos rely on shared components of the UcPS machinery, including p24/TMED family members, GRASP proteins, autophagy factors, and ESCRT-associated components (Noh et al., 2018). However, important distinctions are evident. Whereas UcPS of CFTRΔF508 is primarily driven by TMED3 and GRASP55, these factors were ineffective in promoting proTGFα secretion. This divergence points to cargo-specific deployment of distinct p24-GRASP pairs, suggesting that a conserved core UcPS machinery is adapted by accessory factors to confer substrate specificity.

Functionally, such UcPS pathways may help alleviate ER proteostasis stress by rerouting excess or misfolded proteins that overwhelm the ERAD pathway from degradation toward secretion. TMED9 has also been implicated in promoting colorectal cancer metastasis through a mechanism linked to TGFα signaling (Mishra et al., 2019). TMED9-mediated UcPS of proTGFα may therefore provide a mechanistic basis for this observation. Notably, TMED9 also enhances expression of CNIH4, a cornichon family cargo receptor involved in conventional proTGFα trafficking, suggesting that TMED9 may coordinate multiple secretory routes (Mishra et al., 2019).

### Study limitations

Our conclusions are based on overexpression of proTGFα, which may introduce non-physiological effects, although such conditions may approximate proTGFα-high cancer settings. Activation of the UcPS pathway required elevated TMED9 or GRASP65 levels, consistent with UcPS pathways that are conditionally engaged under stress or high cargo load (Noh et al., 2022). Important limitations include the absence of high-resolution imaging to resolve trafficking intermediates and the need for additional interaction studies to define pathway architecture. Identification of endogenous cargos and physiological triggers will be essential to establish the broader relevance of this UcPS mechanism. Furthermore, testing the contribution of TMED9 and GRASP65 in cancer models with a high proTGFα expression and secretion will be necessary to evaluate its potential disease relevance.

## Methods

### Plasmids

Constructs for FLAG-tagged human proTGFα, proTGFαΔ4 (Wunderle et al., 2016), C-terminally myc-tagged human GRASP55 and GRASP65 as well as the chimeric GRASP mutants (Nüchel et al., 2021), and human RHBDL4 with an N-terminal triple HA-tag (Wunderle et al., 2016) have been described previously. Human TMED9, human TMED7, human TMED3 were amplified from expression vectors containing a signal peptide before a triple FLAG-tag (Knopf et al., 2024) and were subcloned into the same vector with the tag removed. The TMED9-AA mutant (F191A, F192A) was generated using subsequent site-directed mutagenesis.

### Culture of cell lines

HEK293T cells (CRL-3216, ATCC) were grown in DMEM (Gibco) supplemented with 10% fetal bovine serum at 37 °C in 5% CO2. HCT116 cells (Gift from Elmar Schiebel, ZMBH, Heidelberg, DE) were grown in McCoy’s 5A medium (Gibco) supplemented with 10% fetal bovine serum at 37 °C in 5% CO2.

### Transfection

For ectopic expression of proteins, cells were transfected with plasmid DNA 24 h after seeding using 25 kDa linear polyethyleneimine (Polysciences) (Durocher et al., 2002). In a 6-well format, if not stated differently in the figure legends, 500 ng of factors analyzed were transfected. For larger formats the amount of transfected DNA was adjusted according to the surface area. Empty vector DNA was used to keep the amount of transfected DNA constant between all conditions. To knockdown ATG5 and ATG7, cells were transfected with 50 pmol each of ATG5-targeting (5’-TCTCGATATCA-ATGACAGATGACAAAGATG-3‘, 5‘-CTCTCTCGAGTCAATCTGTTGGCTGTGGGATG-3‘) and ATG7 (5’-TATAGAATTCTAATGGCGGCAGCTACGGGG-3‘, 5‘-TATAGCGGCCGC-TCAGATGGTCTCATCATCGC-3‘)-targeting siRNA (Life Technologies) or the same amount of non-targeting siRNA. For knockdown of ALIX, cells were transfected with 50 pmol ALIX-targeting siRNA (ON-TARGETplus SMARTpool human siRNA, Horizon Discovery, NM_013374, 5’-GAAGGAUGCUUUCGAUAAA-3’, 5’-GAACAGAACCUGGAUAAUG-3’, 5’-GAGAGGGUCUGGAGAAUGA-3’, 5’-GCAGUGAGGUUGUAAAUGU-3’) or the same amount of non-targeting siRNA. Cells were treated with 10 µM BB94 24 h before harvesting to inhibit metalloproteases.

### Immunoprecipitation

For crosslinking, cells were incubated for 30 min with 1 mM DSP (Thermo Fisher Scientific) in PBS at room temperature while gently shaking. The crosslinking reaction was quenched by the addition of 20 mM TRIS-HCl pH 7.5 for 15 min. Cells were lysed in solubilization buffer (50 mM HEPES-KOH, pH 7.4, 150 mM NaCl, 2 mM Mg(OAc)2, 10% glycerol, 1 mM EGTA, 10 mM N-ethylmaleimide) supplemented with 1% Triton X-100 (Merck) and EDTA free complete protease inhibitor cocktail (Roche) for 60 min on ice. The Triton-insoluble fraction was pelleted by centrifugation at 16.000 x g for 15 min at 4 °C. The supernatants were diluted 1:1 with lysis buffer without detergent. 240 ng mouse monoclonal anti-myc antibody (clone 9B11) (2276, Cell Signaling Technologies) were added to the samples. Samples were incubated while rotating at 4 °C over-night. Protein G-coated magnetic beads (Thermo Fisher Scientific) were added for 1 h to the samples. Beads were washed according to the manufacturer’s protocol and subsequently eluted with SDS-sample buffer. Samples were analyzed by western blotting.

### Cell Surface Biotinylation

Cells were washed twice with PBS containing 0.9 mM MgCl2, and 0.5 mM CaCl2 before being incubated for 30 min with 0.5 mg/ml Sulfo-NHS-LC-Biotin (Apex-bio) in PBS (+ 0.9 mM MgCl2, 0.5 mM CaCl2) while gently shaking at 4 °C. Afterwards, cells were washed once with PBS supplemented with 0.9 mM MgCl2, 0.5 mM CaCl2, and twice with 100 mM glycine in PBS supplemented with 0.9 mM MgCl2, 0.5 mM CaCl2 to quench the biotinylation reaction. Cells were lysed in lysis buffer (50 mM TRIS-HCL pH 7.5, 0.5% Triton X-100, 150 mM NaCl, 0.1% SDS) with freshy added complete protease inhibitor cocktail (Roche) for 60 min on ice. The detergent-insoluble fraction was pelleted by centrifugation at 16.000 x g for 15 min at 4 °C. 2% of input was taken from the supernatant, before 30 µl pre-washed Streptavidin Sepharose beads (Cytiva) were added to the samples. Samples were incubated while rotating at 4 °C over-night. Beads were washed four times with lysis buffer. Proteins were eluted with 2x SDS sample buffer (50 mM TRIS-pH 6.8, 10 mM EDTA pH 8.0, 5% glycerol, 2% SDS, 0.01% bromophenol blue) containing 50 mM DTT for 30 min at 37 °C. Samples were analyzed by western blotting.

### Antibodies

The following antibodies were used for western blotting with indicated dilutions: mouse monoclonal anti-FLAG M2 (F1804, Sigma Aldrich, 1:1000), mouse monoclonal anti-FLAG-HRP (clone M2, A8592, Sigma Aldrich, 1:1000), rabbit monoclonal anti-myc clone 71D10 (2278, Cell Signaling Technologies, 1:1000), rabbit polyclonal anti-TMED9 (21620-1-AP, Proteintech, 1:1000), rabbit anti-TMED7 (gift from F. Wieland, Heidelberg University, 1:1000), guinea-pig anti-TMED3 (gift from F. Wieland, Heidelberg University, 1:1000), rabbit monoclonal anti-ATP1A1 clone EP1845Y (ab76020, Abcam, 1:5000), rabbit polyclonal anti-ATG5 (A0856, Sigma, 1:500), mouse monoclonal anti-ALIX clone C-11 (sc-271975, Santa Cruz Biotechnology, 1:1000).

### Western blotting

Whole cell lysates were prepared in SDS sample buffer containing 5% 2-mercaptoethanol and incubated for 10 min at 65 °C before separation by Tris-glycine SDS poylacrylamide gel electrophoresis. For samples that were subjected to EndoH and PNGaseF digests, the samples were diluted 1:4 with water and then treated with EndoH (New England Biolabs), PNGaseF (New England Biolabs) or left untreated according to the manufacturer’s instructions before separation by gel electrophoresis. To analyze secreted proteins, cells were cultured 24 h before harvesting in OptiMEM (Thermo Fisher Scientific). Dead cells were removed from the supernatant by centrifugation at 500 x g and 4 °C for 5 min followed by centrifugation at 16.000 x g for 15 min to pellet cell debris. Proteins were precipitated with 10% trichloroacetic acid for 5 min on ice. Precipitated proteins were collected by centrifugation at 16.000 x g and 4 °C for 15 min. Pellets were washed with acetone, dried and solubilized in SDS sample buffer to be analyzed by gel electrophoresis. Where shown, 2,2,2-trichlorethanol (TCE, T54801, Sigma Aldrich) was added to gels. TCE staining was analyzed by ChemiDoc Touch imaging system (Bio-Rad Laboratories) as loading control. Gels were transferred onto PVDF membranes by semi-dry blotting. After incubation with respective antibodies, membranes were analyzed by enhanced chemiluminescence analysis using the ImageQuant 800 Fluor system (Cytiva) with the ImageQuant 800 software (version 2.0.0). Signal intensities were quantified using Fiji (Schindelin et al., 2012).

### Quantification and statistical analysis

If not stated otherwise experiments were conducted in triplicates. Statistical analysis was performed in GraphPad (version 10.2.0).

## Acknowledgments

We acknowledge support by the Center for Molecular Medicine Cologne (CMMC) for funding through the Individual Project Funding Program (Project C10) and the Career Advancement Program to J.N. (Project CAP31). Parts of this work were supported by the Deutsche Forschungsgemeinschaft (DFG, German Research Foundation) through the Research Unit Grant FOR2722 (NU 467/1-1; project No 384170921) to J.N., the DFG Grant NU 467/2-1 (project No 564334266) to J.N.

## Author contribution

Conceptualization: S.S.S., J.N., M.K.L.; Methodology: S.S.S., J.N., M.K.L.; Formal analysis: S.S.S., J.N., M.K.L.; Investigation: S.S.S., I.D.; Writing - original draft: S.S.S., M.K.L.; Writing - review & editing: S.S.S., J.N., M.K.L.; Visualization: S.S.S., M.K.L.; Supervision: S.S.S., M.K.L.; Project administration: M.K.L.; Funding acquisition: J.N., M.K.L.

