## Supplementary material for "A Golgi-bypass secretion mechanism for proTGFα involving TMED9 and GRASP65": Fig. S1 and S2

**Supplementary Information Steigleder *et al.***

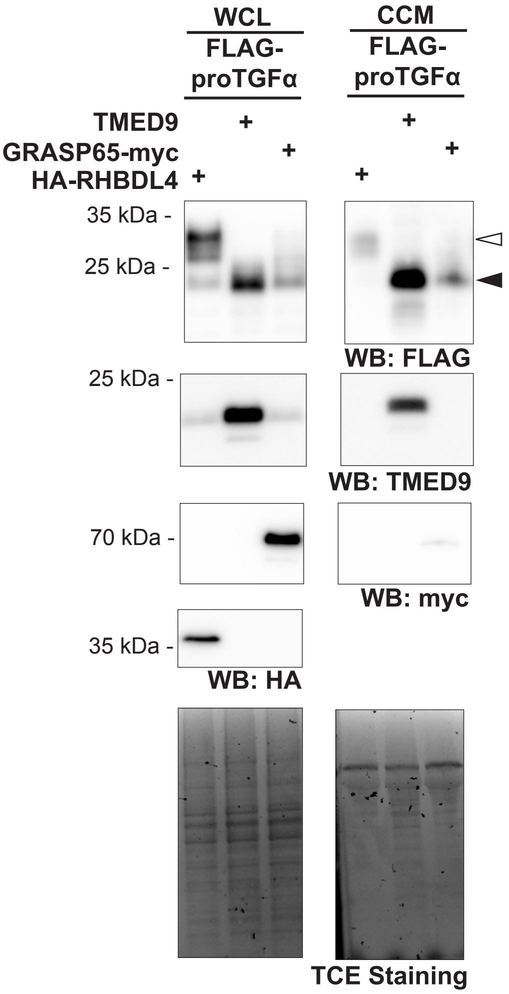


**Fig. S1. Comparison of the effect of ectopic RHBDL4, TMED9 and GRASP65 expression on proTGFα secretion.** Western blot (WB)-based secretion assay using Hek293T co-expressing proTGFα with either HA-RHBDL4, TMED9 or GRASP65-myc. While both TMED9 and GRASP65 expression led to the intracellular accumulation of the ER form of proTGFα, that is also secreted (filled arrow), HA-RHBDL4 expression led to the intracellular accumulation of the Golgi form of proTGFα, that is also secreted (open arrow). TCE staining is used as a loading control. Cells were treated with BB94 (10 µM).

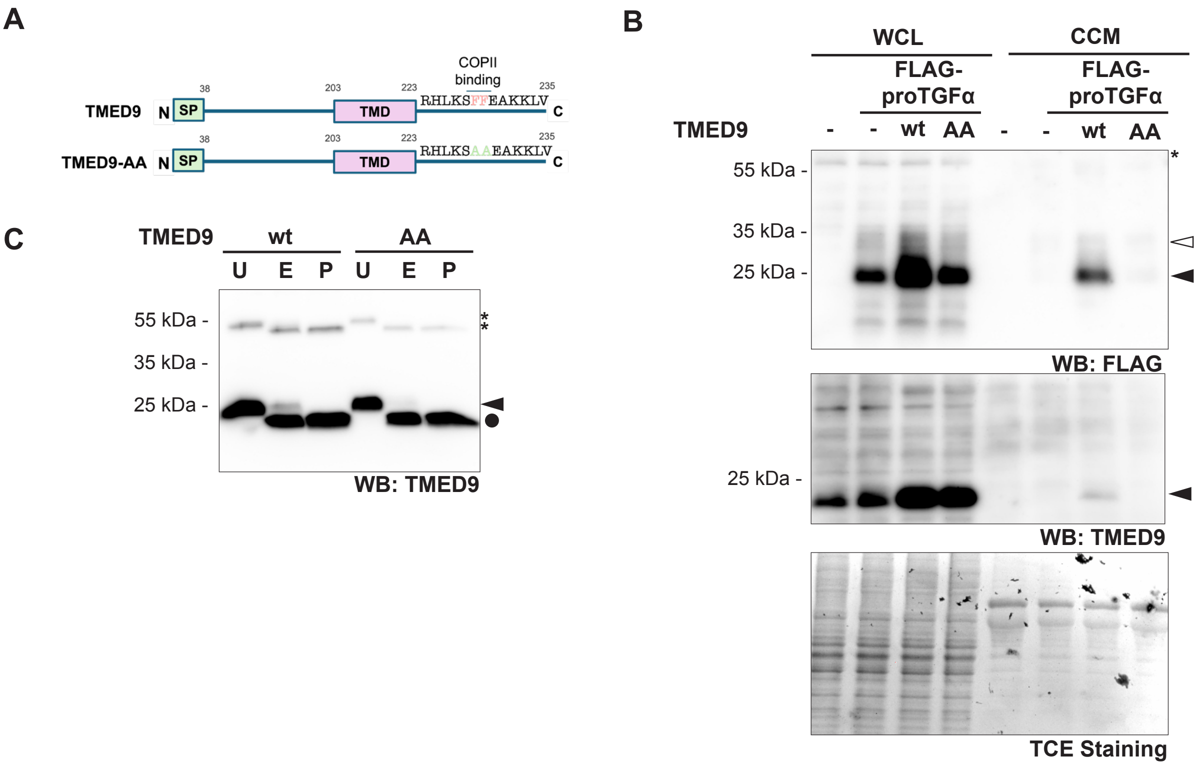


**Fig. S2. TMED9 COPII-binding mutant does not induce UcPS of proTGFα. (A)** Schematic representation of the COPII-binding motif in TMED9 and the mutant (TMED9-AA) where the motif is exchanged to alanine residues. SP, signal peptide, TMD, transmembrane domain. **(B)** Western blot (WB)-based secretion assay using Hek293T cells co-expressing FLAG-proTGFα with either empty vector (-), wild-type TMED9 (wt) or TMED9-AA (AA). Only TMED9 wt induces accumulation and secretion of proTGFα (filled arrow). Unknown bands are indicated with an asterisk. TCE staining is used as a loading control. Cells were treated with BB94 (10 μM). **(C)** WB-analysis following treatment with EndoH (E), PNGaseF (P) or no treatment (U) of Hek293T cells overexpressing either TMED9 wt or AA mutant (filled arrow). Unknown bands are indicated with an asterisk. TMED9-AA is sensitive to EndoH-digest (filled circle) suggesting it is confined to the ER.
